# The rodent hippocampus as a bilateral structure: A review of hemispheric lateralization

**DOI:** 10.1101/150193

**Authors:** Jake T. Jordan

## Abstract

The left and right rodent hippocampi exhibit striking lateralization in some of the very neural substrates considered to be critical for hippocampal cognitive function. Despite this, there is an overwhelming lack of consideration for hemispheric differences in studies of the rodent hippocampus. Asymmetries identified so far suggest that a bilateral model of the hippocampus will be essential for an understanding of this brain region, and perhaps of the brain more widely. Although hypotheses have been proposed to explain how the left and right hippocampi contribute to behavior and cognition, these hypotheses have either been refuted by more recent studies or have been limited in the scope of data they explain. Here, I will first review data on human and rodent hippocampal lateralization. The implications of these data suggest that considering the hippocampus as a bilateral structure with functional lateralization will be critical moving forward in understanding the function and mechanisms of this brain region. In exploring these implications, I will then propose a hypothesis of the hippocampus as a bilateral structure. This discrete-continuous (DC) hypothesis proposes that the left and right hippocampi contribute to spatial memory and navigation in a complementary manner. Specifically, the left hemisphere stores spatial information as discrete, salient locations and that the right hemisphere represents space continuously, contributing to route computation and flexible spatial navigation. Consideration of hippocampal lateralization in designing future studies may provide insight into the function of the hippocampus and resolve debates concerning its function.

## Introduction

Animal experimental models are powerful tools for investigating the cellular and molecular bases of cognition. Non-human animal studies on the hippocampal formation have led to the discovery of spatially selective place cells (O’Keefe & Dostrovsky, 1971), long-term potentiation (Bliss & Lømo, 1973), and sharp wave ripple oscillations (Buzsáki et al., 1992), findings which have provided invaluable insight into the neural bases of spatial cognition and memory. An obstacle in extending findings in animal studies to our understanding of the human hippocampus is the widely held view that the rodent hippocampus does not exhibit the same interhemispheric differences that are seen in humans. Lateralization is an asymmetry, in degree or presence, of a particular neural substrate or process between hemispheres (Concha et al., 2012). The human hippocampus is strongly lateralized with respect to cognitive function (Maguire et al., 1998; O’Keefe et al., 1998; Spiers et al., 2001; Burgess et al., 2002; Maguire & Frith, 2003; Howard et al., 2014). Specifically, the left hippocampus is specialized for episodic, contextual, and long-term autobiographical memory (Spiers et al., 2001; Maguire & Frith, 2003), while the right hippocampus is specialized for navigation (Maguire et al., 1998; Spiers et al., 2001; Howard et al., 2014). However, despite recognition of lateralization in several measures of function, anatomy and physiology (Kawakami et al., 2003; Shinohara et al., 2008; Kohl et al., 2011; Shipton et al., 2014; Benito et al., 2016; Villalobos et al., 2017), a vast majority of studies on the rodent hippocampus do not take lateralization into account.

Historically, rodent hippocampal lateralization has been ignored because the rodent hippocampus has substantial bilateral projections between hippocampal subfields, which are thought to be absent or considerably weaker in primates (Wilson et al., 1987; Amaral & Lavenex, 2007). It has been hypothesized that a lack of interhemispheric communication between the hippocampi in humans has led to functional lateralization in this species (Zaidel, 1995). Supporting this idea, navigation in the Morris Water Maze (MWM), a commonly used behavioral test to assess rodent hippocampal function, relies on both hippocampi for optimal performance in rats (Fenton & Bures, 1993). Thus, it is widely thought that the rodent hippocampus is not functionally lateralized. However, several studies have indicated that the human brain may indeed have direct interhemispheric projections between the left and right hippocampi (Gloor et al., 1993; Rosenzweig et al., 2011; Lacuey et al., 2015). Further, hippocampal lateralization has been reported in function (Klur et al., 2009), physiology (Benito et al., 2016) and chemistry in rats (Louilot & Le Moal, 1994; Wolff et al., 2008), and in function (Shinohara et al., 2012, Shipton et al., 2014; El-Gaby et al., 2016), synaptic plasticity (Kohl et al., 2011; Shipton et al., 2014) and anatomy in mice (Kawakami et al., 2003; Shinohara et al., 2008; Shinohara & Hirase, 2009). As discussed below, some of the most intriguing hemispheric asymmetries in rodent hippocampal physiology are not easily identifiable using classical experimental techniques. These asymmetries were only recently uncovered using advanced methods. Importantly, a recent study showed that failure to consider differences between the left and right rodent hippocampi in cellular quantification experiments can lead to incomplete and inconsistent conclusions (Jordan et al., 2019). Collectively, these findings indicate the importance of understanding the hippocampus as a bilateral structure and of taking into account differences across hemispheres.

Here, I will review literature on hippocampal lateralization in rats, mice and humans and the relation of these data to memory and spatial cognition. I will first briefly review the genetics of body asymmetry and how these genes may influence cognitive function. I will then describe major findings on hippocampal lateralization discovered using rodent models (Table 1), the methods leading to these discoveries, and why lateralization is difficult to detect using classic experimental protocols. I will argue that hemispheric asymmetries in functional specialization in rodents may be more similar to that seen in humans than traditionally thought, and that rodent models may lead to novel insights regarding interhemispheric contributions to hippocampal function. Finally, I will review theories of the cognitive contributions of the left and right hippocampi and propose a novel hypothesis for how the left and right hippocampi process spatial information and contribute to hippocampal function.

**Table 1.1:**
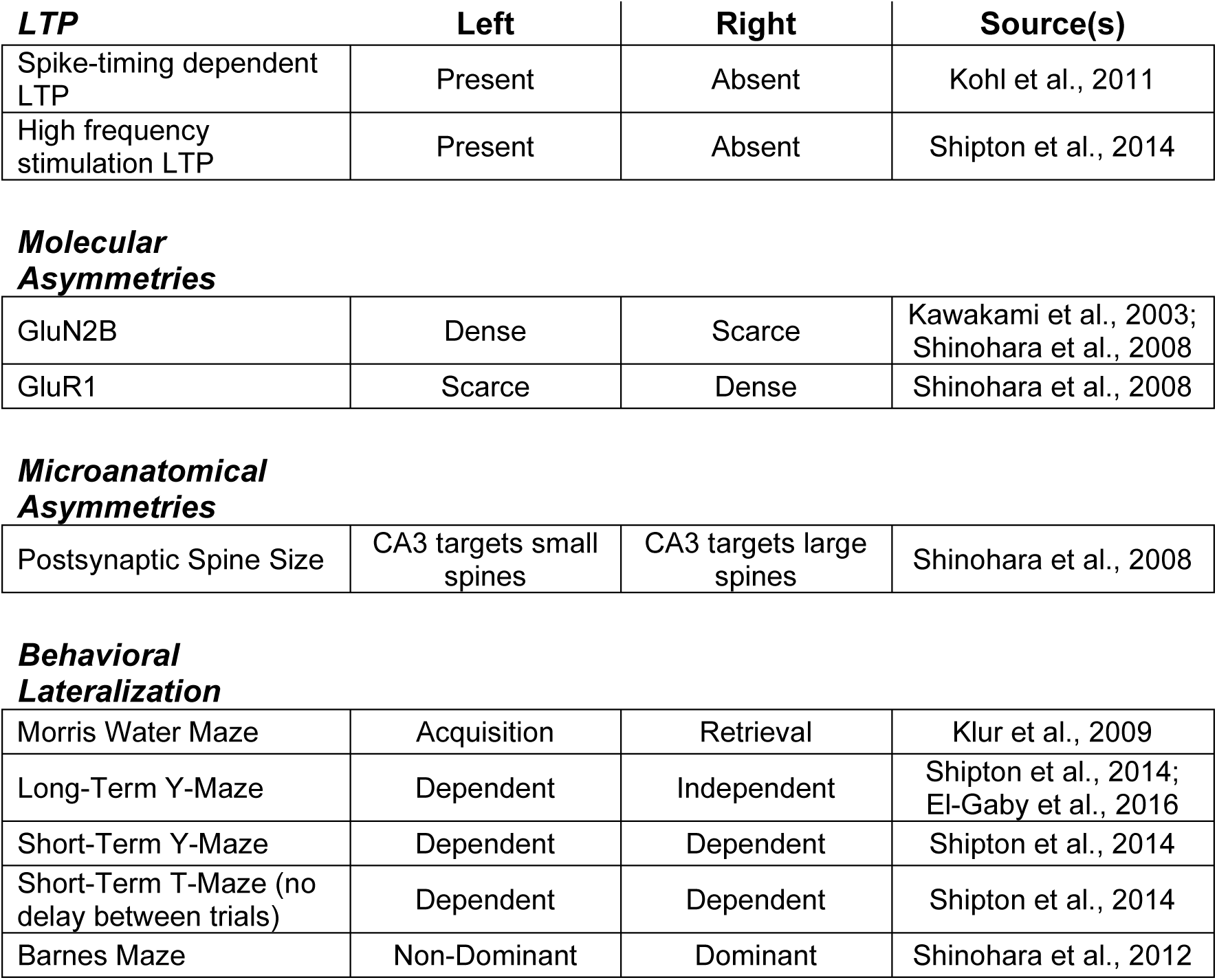
Summary results of rodent hippocampal lateralization studies

| ***LTP*** | **Left** | **Right** | **Source(s)** |
| --- | --- | --- | --- |
| Spike-timing dependent LTP | Present | Absent | Kohl et al., 2011 |
| High frequency stimulation LTP | Present | Absent | Shipton et al., 2014 |
| ***Molecular Asymmetries*** |  |  |  |
| GluN2B | Dense | Scarce | Kawakami et al., 2003;<br>Shinohara et al., 2008 |
| GluR1 | Scarce | Dense | Shinohara et al., 2008 |
| ***Microanatomical Asymmetries*** |  |  |  |
| Postsynaptic Spine Size | CA3 targets small spines | CA3 targets large spines | Shinohara et al., 2008 |
| ***Behavioral Lateralization*** |  |  |  |
| Morris Water Maze | Acquisition | Retrieval | Klur et al., 2009 |
| Long-Term Y-Maze | Dependent | Independent | Shipton et al., 2014;<br>El-Gaby et al., 2016 |
| Short-Term Y-Maze | Dependent | Dependent | Shipton et al., 2014 |
| Short-Term T-Maze (no delay between trials) | Dependent | Dependent | Shipton et al., 2014 |
| Barnes Maze | Non-Dominant | Dominant | Shinohara et al., 2012 |

## Conservation of Organismal and Brain Lateralization

Cerebral lateralization was once considered to be a uniquely human phenomenon, perhaps enabling behaviors or cognitive functions that are unique to humans. Since the 19^th^ century, it has been known that the human brain is lateralized with respect to verbal communication and comprehension. Lateralization in non-human animals was not reported until long after the initial reports on humans. Lateralization with respect to various forms of memory has since been reported not only in humans (e.g. Spiers et al., 2001), but in macaques (visual memory in *Macaca nemestrina*; Doty et al., 1999), fruit flies (olfactory cued fear memory in *Drosophila melanogaster*; Pascual et al., 2004), honeybees (olfactory cued reward memory in *Apis mellifera*; Rogers & Vallortigara, 2008), snails (food aversion memory in *Helix lucorum*; Kharchenko et al., 2010) and zebra finches (conspecific song memory in *Taeniopygia guttata*; Tsoi et al., 2014). Further, hippocampal lateralization of spatial memory in particular has also been reported in humans (Spiers et al., 2001) as well as homing pigeons (*Columba livia*; Kahn & Bingman, 2004), mice (*Mus musculus*; Shinohara et al., 2012; Shipton et al., 2014; El-Gaby et al., 2016) and rats (*Rattus norvegicus*; Klur et al., 2009). In fact, flies and mice with asymmetric neural circuitry have superior memory compared to those with symmetric brains (flies: Pascual et al., 2004; mice: Kawakami et al., 2008; Goto et al., 2010) and the degree of left-dominance in the songbird auditory memory system positively correlates with song memory (Tsoi et al., 2014). These data indicate that lateralization in the neural systems underlying various forms of memory may be more of a rule across animals than a set of exceptions.

### Genetics of body and neural lateralization

How does lateralization happen in the first place? The genetics of organismal lateralization may be conserved in invertebrates and in vertebrates (Blum & Ott, 2018; Yuan & Brueckner, 2018). In mammals, thoracic and abdominal organs have a ubiquitous asymmetric arrangement and the development of this lateralization is dependent upon leftward flow of growth factors generated by the chiral beating of cilia during embryonic development (Nonaka et al., 2002). Loss of ciliary function in humans leads to a condition called primary ciliary dyskinesia (PCD). In patients with PCD, among other symptoms is a randomized laterality of thoracic and abdominal organs (Lobo et al., 2014), which is seen in other organisms with ciliary dysfunction (Nonaka et al., 2002). Artificially induced rightward nodal flow reverses organ laterality in wild-type mice and can induce stereotyped laterality in mice with immotile cilia that typically have a randomized laterality (Nonaka et al., 2002). A number of genes have been implicated in PCD and in organ lateralization (Lobo et al., 2014). In rodents, multiple genes appear to affect not only organ lateralization but also spatial memory and hippocampal lateralization. *DYX1C1* is associated with PCD in humans and deletion of this gene in mice produces a PCD-like phenotype, including altered organ laterality (Tarkar et al., 2013). In rats, *in utero* administration of *DYX1C1* RNAi impairs spatial working memory (Szalkowski et al., 2011). In addition, mutations to the gene *left-right dynein* randomize organ lateralization and abolish hippocampal lateralization in mice (Kawakami et al., 2008).

Interestingly, these mutant mice exhibit deficits in spatial reference and working memory (Goto et al., 2010). Collectively, these data indicate that the genes involved in determining thoracic and abdominal lateralization during development may be conserved in mammals and may also play a role in the establishment of functional nervous system asymmetries, including in the mammalian hippocampus (Figure 1). It is not known whether patients with PCD or other conditions in which organ laterality is affected have altered hippocampal lateralization and there have been no reports that these patients have altered hippocampal memory function.

**Figure 1.**
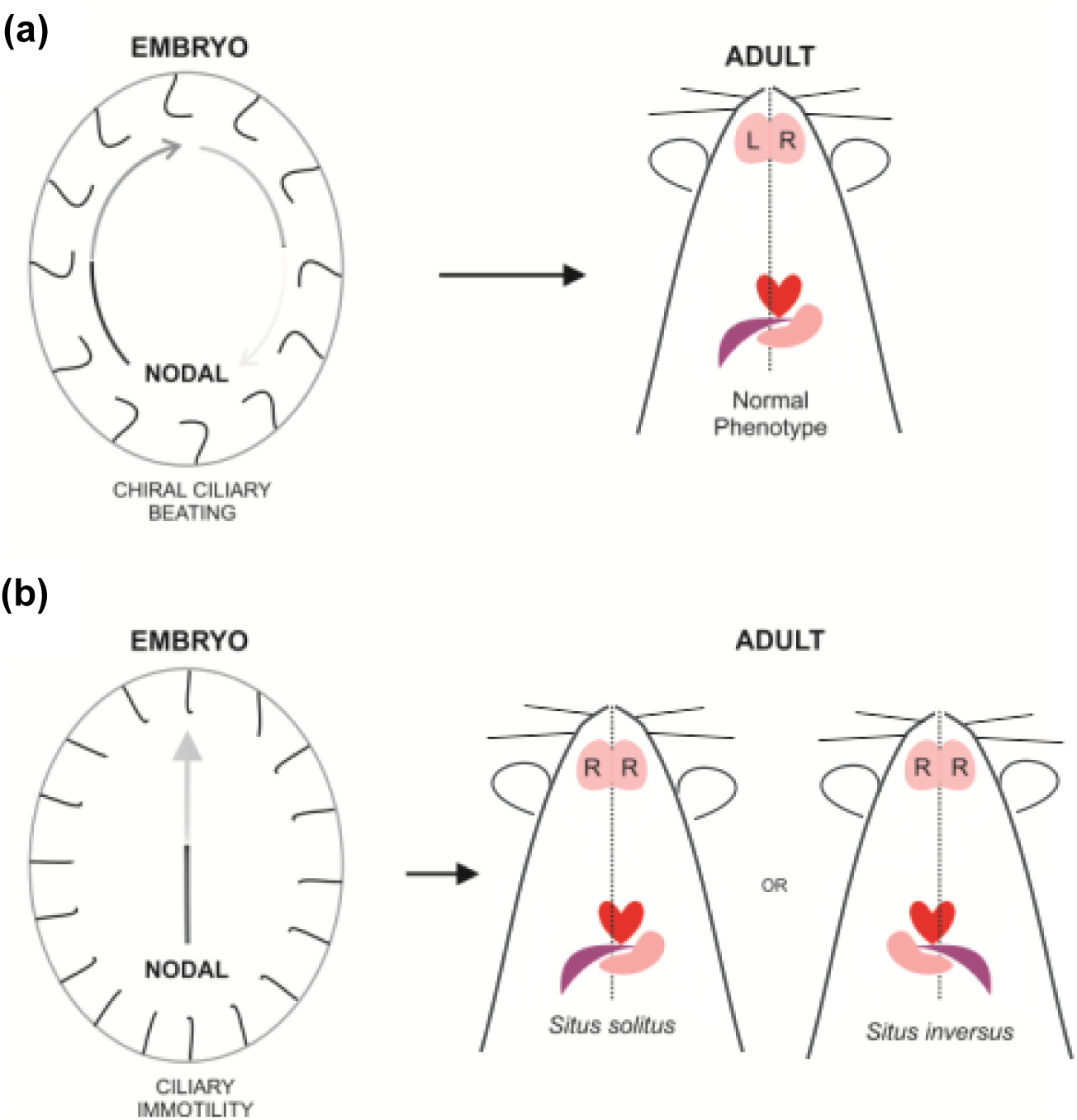
Lateralization of organ systems and of neural memory systems in mammals. (a) The chiral beating of cilia produces a leftward nodal flow in the developing embryo. As adults, mammals have a stereotyped laterality of thoracic and abdominal organs (Nonaka, et al., 2002). Wild type mice with normal ciliary function have an asymmetric hippocampus (Kawakami et al., 2003; Shinohara et al., 2008; Kawakami et al., 2008). (b) Loss of ciliary motility results in a randomized organ laterality in mice and humans (Nonaka et al., 2002; Kawakami et al., 2008; Lobo et al., 2014). Affected mice have a bilateral right isomerism of hippocampal circuitry, regardless of direction of organ laterality (Kawakami et al., 2008). Hippocampal lateralization has not yet been studied in humans with ciliary immotility.

## Lateralization at Mouse CA3/CA1 Synapses

Area CA1 of the mouse hippocampus receives excitatory input from both the ipsilateral (via Schaffer collateral projections) and contralateral (via commissural projections) CA3 (Figure 2a), often referred to as the “Schaffer collateral-commissural” pathway. Commissural fibers form the ventral hippocampal commissure (VHC). It should be pointed out that this term is not in relation to the functional distinction between the dorsal and ventral hippocampus (Kjelstrup et al., 2002) as fibers within the VHC innervate the entire dorso-ventral extent of the contralateral hippocampus (Amaral & Lavenex, 2007). Manipulations that impair LTP at CA3-CA1 synapses also impair long-term memory (Wong et al., 1999). As will be discussed in this section, intriguing hemispheric asymmetries have been reported in mice in molecular anatomy and physiology at CA3-CA1 synapses depending on the hemispheric origin of CA3 input. That is, projections originating from left CA3 that target both left and right CA1 dendritic spines differ anatomically and physiologically from projections that originate from right CA3 that target left and right CA1 spines.

**Figure 2.**
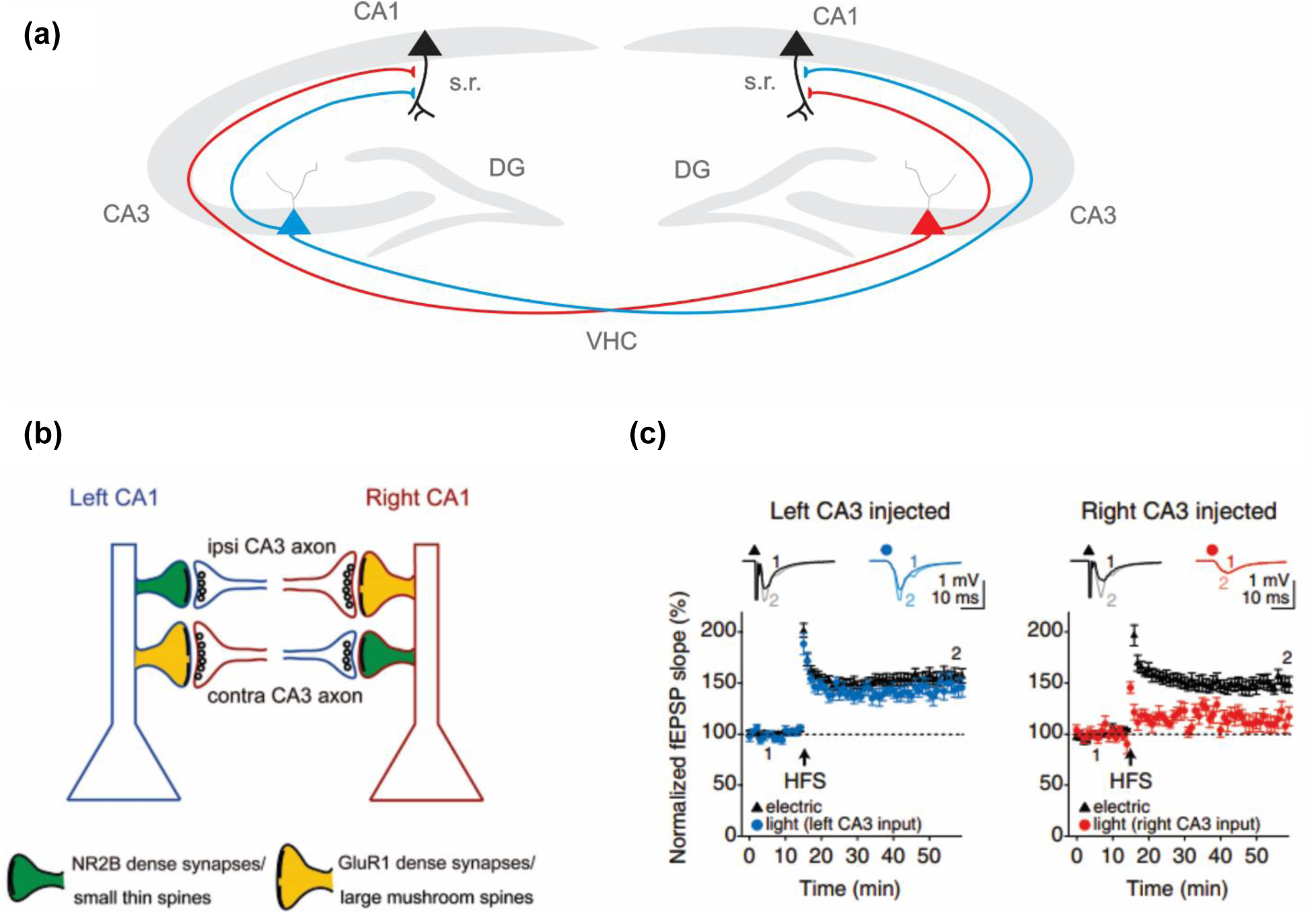
Lateralization of synaptic plasticity at Schaffer collateral-commissural synapses depends on CA3. (a) Left and right CA3 (blue and red, respectively) project to ipsilateral and contralateral CA1. (b) Synapses with CA1 neurons show different molecular composition and microanatomical properties depending on whether they receive input from left (blue axons) or right (red axons) CA3, regardless of which hemisphere the postsynaptic target resides in. Specifically, left and right CA1 spines receiving left CA3 input are smaller in PSD area and volume than those receiving right CA3 input. CA1 spines receiving left CA3 input have high densities of GluN2B (NR2B in figure), whereas those receiving right CA3 input have high densities of GluA1 (GluR1 in figure) (from Shinohara et al., 2008, *PNAS*, Copyright (2008) National Academy of Sciences). (c) LTP in CA1 after high frequency stimulation (HFS) of left or right CA3 (arrow). In this experiment, HFS was delivered via an electrical stimulus. To examine CA1 postsynaptic responses to CA3 input, test pulses were administered to CA3 using electrical or optical stimuli. Test pulses using electrical stimuli (black) revealed robust potentiation but no lateralization. However, selective optogenetic stimulation of left CA3 neurons (blue) reveals LTP in left and right CA1, while optogenetic stimulation of right CA3 reveals very low potentiation of left and right CA1 (from Shipton et al., 2014, *PNAS*). DG, dentate gyrus; s.r., stratum radiatum; VHC, ventral hippocampal commissure.

### Asymmetry of postsynaptic receptor subunits at Schaffer collateral-commissural synapses

CA3-CA1 synapses differ substantially in microanatomy, molecular composition, and capacity for LTP depending on the hemispheric origin of CA3 input and independent of which hemisphere the postsynaptic CA1 spine resides in. In a landmark study, VHC transection, the severing of all commissural fibers projecting from CA3 to the contralateral CA1, identified intriguing molecular asymmetries of the mouse hippocampus (Kawakami et al., 2003). Following surgery, the left and right CA1 of VHC-transected mice received only ipsilateral input from left and right CA3, respectively. Both the left and right CA1 of surgically naïve control mice received input from both left and right CA3. In VHC transected mice, the authors identified a higher density of the GluN2B (also referred to as ε2 or NR2B) protein in *stratum radiatum* of left CA1 than right CA1. Lateralization was not seen in surgically naïve mice with an intact VHC, suggesting that this lateralization was due to the hemispheric origin of CA3 input. The authors performed whole-cell slice recordings and measured NMDA EPSCs in CA1 apical dendrites following stimulation of the *stratum radiatum* and found that neurons receiving left CA3 input showed a greater sensitivity to GluN2B antagonism (via bath application of the GluN2B antagonist Ro-25-6981) than those receiving right CA3 input, but only in VHC-transected mice. Interestingly, the authors found the reverse pattern of lateralization with a higher density of GluN2B expression in the right *stratum oriens* than in the left in VHC-transected mice. Further, EPSCs in the right CA1 showed a greater sensitivity to GluN2B blockade following stimulation of the *stratum oriens*. It is not clear what the functional significance of this reverse pattern of lateralization in the *stratum radiatum* and *oriens* is. However, a majority of CA1 synapses lie in *stratum radiatum* and subsequent studies of physiological lateralization examined the properties of the Schaffer collateral-commissural pathway in *stratum radiatum*. A follow-up study determined that asymmetric GluN2B density in CA1 was specific to pyramidal neurons as interneurons showed no such laterality (Wu et al., 2005). As GluN2B is closely associated with the capacity for LTP (Lisman et al., 2002), it appeared that hippocampal LTP may demonstrate interhemispheric asymmetries. However, experimental limitations (i.e. inability to selectively stimulate unilateral CA3 fibers) prevented the authors from testing this possibility directly. Despite these limitations, Kawakami et al., (2003) were the first to suggest that physiological processes thought to be essential for hippocampal function (e.g. LTP) may be asymmetric across hemispheres.

A follow-up study confirmed the lateralization of GluN2B density and further characterized bilateral expression of other glutamate receptor subunits and postsynaptic targets of left and right CA3 axons (Shinohara et al., 2008). GluA1 (also referred to as GluR1), an AMPA receptor subunit associated with LTP saturation, was shown to have higher expression levels in both left and right CA1 synapses that receive input from the right CA3 (Shinohara et al., 2008), reflecting a mirror asymmetry to that seen in GluN2B. There was no observed asymmetry in GluN2A, GluA2, or GluA3, and the functional implications of this pattern are unclear. Additionally, the authors characterized the morphology of spines in CA1 synapsing with left or right CA3 axons. To do this, they injected a viral vector to drive expression of axonal GFP into either the left or right CA3 of intact mice. These axons were then traced via serial electron microscopy to their dendritic targets in CA1. These dendritic spines were then digitally reconstructed allowing for volumetric quantification of spine heads. They found that bilateral CA1 spines targeted by right CA3 axons had a larger volume and synaptic surface area than CA1 spines targeted by left CA3 axons. Further, a greater proportion of right CA3-targeted CA1 spines displayed the larger mushroom head phenotype than spines with a thin phenotype. This study concluded with a model of hippocampal circuitry in which left CA3 fibers bilaterally targeted small, GluN2B-rich CA1 spines while right CA3 fibers bilaterally targeted large GluA1-rich spines (Figure 2b).

The metabotropic glutamate receptor subunit mGluR5 is associated with long-term depression (LTD) at hippocampal Schaffer collateral-commissural synapses (Kirschstein et al., 2007). Interestingly, this subunit shows a similar pattern of lateralization as GluN2B, being more densely expressed at CA1 synapses receiving left CA3 input (Shinohara & Hirase, 2009). To summarize, GluN2B and mGluR5, molecules associated with complementary forms of synaptic plasticity (Lisman et al., 2002; Kirschstein et al., 2007) are expressed at higher densities at CA1 synapses receiving left CA3 input (Kawakami et al., 2003; Shinohara et al., 2008; Shinohara & Hirase, 2009). GluA1, a molecule associated with saturation of LTP and synaptic maturity (Shinohara et al., 2008), is dominant at CA1 synapses receiving right CA3 input (Shinohara et al., 2008). These studies established the presence of molecular hemispheric lateralization in hippocampal neural circuitry.

### Synaptic physiology at Schaffer collateral-commissural synapses

An early idea regarding how the hippocampus may acquire and temporarily store new memories is the synaptic memory hypothesis. This hypothesis suggests that memory may be stored via activity-dependent changes in the strength of synaptic transmission (Bliss & Collingridge, 1993). Though recent data has called this idea into question (Chen et al., 2014; Ryan et al., 2015), it is clear that synaptic plasticity and memory are intimately linked. LTP is a long-lasting form of synaptic plasticity in which synaptic transmission is enhanced on the timescale of hours to days following a high frequency train of presynaptic action potentials (Bliss & Lømo, 1973), or a presynaptic action potential during a time of postsynaptic depolarization (Levy & Steward, 1983; Gustafsson & Wigström, 1986). LTP at Schaffer collateral-commissural synapses between CA3 and CA1 is associated with hippocampus-dependent memory formation (Wong et al., 1999).

As the density of GluN2B expression depends on the hemisphere of origin of CA3 projections (Kawakami et al., 2003), and as GluN2B is associated with the potential for LTP induction (Lisman et al., 2002), these findings led to the hypothesis that LTP at Schaffer collateral-commissural synapses may be left-lateralized (Kohl et al., 2011). However, because CA1 neurons form synapses with presynaptic terminals from both ipsilateral and contralateral CA3, electrically stimulating Schaffer collateral-commissural fibers masks potential lateralization of the terminals originating in the left or right CA3 (Kawakami et al., 2003; Kohl et al., 2011; Shipton et al., 2014). To overcome this limitation, the left or right CA3 was injected with a viral vector carrying the gene for channelrhodopsin-2, a light-gated cation channel (Kohl et al., 2011). This ensured that the fibers originating from CA3 in only one hemisphere could be selectively stimulated using an optical stimulus in a slice preparation, as indicated by a lack of cross-facilitation between optical and electrical stimuli. t-LTP, a form of LTP in which presynaptic stimulation is followed closely by a train of postsynaptic action potentials, was performed on slices from either hemisphere. Presynaptic optical stimulation of only the left CA3 resulted in t-LTP at Schaffer collateral-commissural synapses in either left or right CA1. However, presynaptic stimulation of right CA3 produced no t-LTP in either left or right CA1. Moreover, electrical stimulation of CA3 in either hemisphere induced t-LTP at Schaffer collateral-commissural synapses, confirming that the asymmetry had been masked using traditional methodologies (as in Kohl et al., 2011; Shipton et al., 2014).

The authors found that NMDA receptor currents and NMDA:AMPA receptor ratios were not lateralized. However, GluN2B subunit-selective NMDA receptor antagonists blocked postsynaptic NMDA currents at Schaffer collateral-commissural synapses more during left CA3 stimulation than during right CA3 stimulation. This supported previous studies finding a higher GluN2B density in synapses originating from the left CA3 (Kawakami et al., 2003; Shinohara et al., 2008). Further, the lack of asymmetry in NMDA receptor currents indicates that the lateralized capacity for t-LTP induction was due to postsynaptic GluN2B expression, and likely not a consequence of asymmetric NMDA receptor expression or presynaptic projection strengths.

Using a similar optogenetic approach, Shipton et al. (2014) examined whether high frequency stimulus (HFS) LTP, which is not affected by GluN2B antagonism, is also lateralized. To do this, they injected left or right CA3 with a ChR2-containing virus (as in Kohl et al., 2011). Since optical stimulation cannot produce a response frequency comparable to standard HFS induction protocols, an electrical stimulus of 100 Hz was used for HFS. Potentiation was tested using optical pulses to left or right CA3 while recording field potentials in CA1. Optical stimuli revealed potentiation of left-CA3 inputs to CA1 but not of right-CA3 inputs (Figure 2c). This effect was again masked when using electrical stimuli to elicit postsynaptic responses after HFS. Thus, both t-LTP and HFS-induced LTP have been demonstrated to be left-dominant at Schaffer collateral-commissural synapses (Kohl et al., 2011; Shipton et al., 2014), indicating a left hemisphere dominance of LTP.

Is left-lateralized plasticity a rule at Schaffer collateral-commissural synapses? So far, only lateralization of LTP at Schaffer collateral-commissural synapses has been demonstrated and only in mice (Kohl et al., 2011; Shipton et al., 2014). Although one study found a lack of asymmetry of Schaffer collateral-commissural LTD in the hippocampus of wild-type mice, this study used an NMDA receptor-dependent induction protocol (O’Riordan et al., 2018). As there is no hemispheric asymmetry of NMDA receptor currents or NMDA:AMPA ratio (Kohl et al., 2011), this result is not surprising. As mGluR5 is associated with LTD and has lateralized expression similar to GluN2B (Shinohara & Hirase, 2009), it is possible that optical LTD induction using a mGluR5-dependent protocol may reveal an asymmetry of this form of plasticity. Further, a recent study found equivalent LTP following electrical stimulation of the left and right CA3 in rats (Martin et al., 2019). This is not surprising as previous studies in mice found that electrical stimuli recruit multiple pathways and can mask hemispheric asymmetries in LTP (Kohl et al., 2011; Shipton et al., 2014). Nonetheless, these findings suggest a need for studies of molecular and physiological lateralization in other species in addition to the mouse.

To date, it appears that LTP at Schaffer collateral-commissural synapses only occurs at CA1 synapses receiving left CA3 input (Figure 2). Considering the relationship between LTP and hippocampal function, these findings have profound implications for our understanding of the hippocampus and call into question the viability of approaching the hippocampus as a bilaterally homogenous structure.

## Functional Lateralization in Humans, Rats and Mice

### Spatial memory in humans

In humans, both the left and right hippocampi appear to be active during goal-oriented spatial navigation but may play different roles in such tasks (Maguire et al., 1998; O’Keefe et al., 1998; Spiers et al., 2001; Burgess et al., 2002; Howard et al., 2014). PET imaging of healthy human subjects found that both the left and right hippocampi demonstrated increased metabolic activity during goal-oriented virtual navigation (Maguire et al., 1998). However, activity in only the right hippocampus positively correlated with navigation accuracy. Interestingly, activity in only the right caudate nucleus significantly correlated with virtual navigation speed. Thus, the authors hypothesized that the right hemisphere may be specialized for spatial navigation. In a subsequent study of patients with unilateral medial temporal lobe (MTL) lesions, subjects explored a virtual town that consisted of multiple rooms where different characters could give them a particular object. Patients with lesions to the left MTL were impaired for contextual aspects of episodic memories, such as identifying the room in which they saw a character (Spiers et al., 2001). In this same study, subjects were made to navigate through the virtual town to a particular location. An image of this location was present to the subject throughout the navigation process. Despite a constant display of the goal location, patients with right MTL lesions were less accurate in navigating to the goal than left-lesion patients or controls. fMRI imaging of BOLD activity during spatial navigation has revealed insight into how the right hippocampus may perform its specialized function of spatial navigation (Howard et al., 2014). In this study, right MTL structures such as the hippocampus and entorhinal cortex, appeared to signal distance to and direction of the goal location. These distance and direction signals were not seen in the left hemisphere.

Considering how patients with left or right MTL damage may function in a more natural setting, it appears that goal-directed spatial navigation would be impaired by lesions to either hemisphere, as left hemisphere damage would impair memory for particular places and right hemisphere damage would impair the ability to navigate to those places. Because an image of a goal location would not be constantly present as it was in the virtual task, left-lesioned subjects would forget which goal location they would be searching for. Right-lesioned subjects may remember perfectly well which location they are searching for but cannot formulate a route to get there. These data demonstrate the requirement of both hemispheres for goal-directed navigation, consistent with clinical studies, and suggest that tasks that require both memory for particular locations and the ability to navigate to them may be insufficient to resolve functional specialization across hemisphere (Maguire et al., 1998; O’Keefe et al., 1998; Spiers et al., 2001; Burgess et al., 2002; Howard et al., 2014).

### Long-term spatial memory in rats

Inactivation of either the left or right hippocampus throughout training and testing equally impair water maze performance in rats (Fenton & Bures, 1993). However, given that this study performed inactivation manipulations throughout the experiment, it was not possible to resolve whether the left and right hippocampi differentially contribute to acquisition and retrieval. In an elegant study, the timing of unilateral hippocampal inactivation via lidocaine infusion was found to affect retrieval in a spatial water maze task, depending on which hemisphere was inhibited and when (Klur et al., 2009). In their first series of experiments, rats were tested on their ability to retrieve the location of a well-learned escape platform: After 6 days of drug-free training, a probe trial was conducted during which the escape platform was removed from the pool and either the left, right, both, or neither hippocampus was inactivated prior to the probe trial, thus testing the ability to navigate to a learned location.

Inactivation of either the right or both hippocampi impaired selective searching for the escape platform (measured by duration spent in the correct quadrant), whereas left hippocampus inactivation had no effect. Therefore, after 6 days of learning, only the right hippocampus was necessary to locate the platform on a learned water maze. In a parallel series of experiments, the authors examined how hippocampal inactivation during acquisition impaired later recall of the escape platform location during a drug-free probe trial. If either the left or both hippocampi were inactivated during the 6 days of training, performance on the drug-free probe trial was impaired. However, inactivation of the right hippocampus during training did not have an effect. Thus, the left hippocampus was needed for acquisition of a spatial memory but was not needed for retrieval of a well-learned spatial memory. Conversely, the right hippocampus was required for expression of a spatial memory but was not required during acquisition to store spatial memories. To explain these data, the authors proposed a hypothesis in which spatial memories are acquired by the left hippocampus and then transferred to the right hippocampus for storage and retrieval, although subsequent data has since called this hypothesis into question (Shipton et al., 2014; El-Gaby et al., 2016).

### Spatial memory in mice

Spatial memory abilities of the left and right hemisphere have also been tested by severing the VHC and corpus callosum in mice and then stitching the left or right eye shut (Shinohara et al., 2012). In mice, 3-5% of retinal output axons project ipsilaterally as opposed to 50% in humans (Erskine & Herrera, 2014). With an intact optic chiasm, a majority of visual information was input to the hemisphere contralateral to the intact eye and this information did not subsequently cross hemispheres. Thus, left eye-intact mice processed a majority of visual input in the right hemisphere and vice versa for right eye-intact mice. Mice were trained over 15 days on a Barnes Maze task in which they had to use extramaze spatial cues to locate an escape hole. There was no difference between left and right eye-intact mice during acquisition. Performance on a probe trial was better in mice that used their right hippocampus than those that used primarily their left hippocampus as indicated by more time spent searching near the learned escape hole. Notably, mice using their left hippocampus still searched selectively near the escape hole, just not as selectively as mice using their right hippocampus. Follow-up experiments showed that both hippocampi were capable of spatial processing as there was no difference between the two groups on a spatial T-Maze task. Finally, there was no difference between groups following contextual fear conditioning, though it is not clear whether sensory modalities other than visual input may have played a role in contextual recall. These data were interpreted as a right-dominance of spatial memory.

However, there are several caveats to be considered when interpreting this study. First, although a majority of visual input was indeed processed by one hemisphere, there is still information received by each eye that is processed by the non-dominant hemisphere. Second, the left and right hippocampi were not inactivated and thus may still have contributed to spatial processing despite reduced visual input, perhaps being driven by peripheral sensorimotor input via the path integration system (McNaughton et al., 1996, 2006). Finally, it is not clear if functional asymmetries were the result of lateralized hippocampal processing or lateralized processing in other brain regions.

In a study of functional hippocampal lateralization in rats, short-term spatial memory was found to be dependent on both hippocampi while long-term spatial memory acquisition was found to be dependent on the left hippocampus only, consistent with left-lateralization of water maze acquisition (Klur et al., 2009) and in contrast to a proposed right-lateralization of spatial memory (Shinohara et al., 2012). The contributions of left and right CA3 to long-term spatial memory in mice were examined using unilateral optogenetic inhibition *in vivo*. Left or right CA3 were inhibited during a long-term Y-Maze task. In this paradigm, one arm of a three-arm Y-Maze is rewarded and mice are placed in one of the other two arms to start a training trial. Mice must then use the configuration of distal spatial cues to determine which arm is rewarded.

Inhibition of left CA3 impaired acquisition of this task even after 11 days of training, however, right CA3 inhibition had no effect. A follow-up study confirmed this finding, showing that optogenetic silencing of left CA3 axons in CA1 impaired spatial Y-Maze acquisition over 10 days of training (El-Gaby et al., 2016). Again, inactivation of right CA3 axons had no effect. These data, indicating a lack of necessity of right CA3 in the long-term Y-Maze task, dispute previous hypotheses of hippocampal lateralization, such as the interhemispheric transfer of spatial engrams (Klur et al., 2009) and right-dominance of spatial memory (Shinohara et al., 2012).

Thus, to explain these data, the authors suggested a time-dependent lateralization of spatial memory in which both hemispheres initially contribute, but over time, learning in the left hippocampus via activity-dependent synaptic plasticity lessens the need for the right hippocampus (Shipton et al., 2014; El-Gaby et al., 2015). However, this hypothesis does not explain the necessity of the right hippocampus in memory retrieval on a well-learned water maze task (Klur et al., 2009), indicating the need for a new hypothesis of bilateral hippocampal function (discussed below).

A partial summary of the consistent findings across humans, rats and mice suggests that both hippocampi contribute to spatial information processing (Maguire et al., 1998; Spiers et al., 2001; Klur et al., 2009; Shinohara et al., 2012; Shipton et al., 2014). With respect to left hippocampal function, it appears that in rats and mice, the left hippocampus is needed for acquisition of long-term spatial memory engrams during learning (Klur et al., 2009; Shipton et al., 2014; El-Gaby et al., 2016). There was no lateralization of acquisition in split-brain mice with laterally-biased visual processing (Shinohara et al., 2012), however, neither hippocampus was inhibited in this experiment. In humans, damage to the left MTL impairs spatial memory for particular places (Spiers et al., 2001). It is not yet clear if left hippocampal data in rodents and in humans reflect similar cognitive functioning. The right hippocampus appears to be required for spatial navigation in a well-learned environment in rats (Klur et al., 2009) and in humans (Spiers et al., 2001) but may also contribute to short-term spatial memory in mice (Shipton et al., 2014). It is not yet clear if these data are a function of a particular cognitive process facilitated by the right hippocampus that has not yet been identified.

### Comparison of human and rodent place memory and navigation studies

Given the divergent effects of left and right MTL damage in humans on performance of spatial memory tasks (e.g., Spiers et al., 2001), it appears that the phenomenon commonly referred to as “spatial memory” may be broken down into separate cognitive subprocesses. In one process, memories for salient locations and events occurring at these locations are acquired, stored and retrieved. This process appears to be facilitated by the left hippocampus. In another process, navigational routes between salient locations, such as a start and a goal location, are computed and then followed. This process appears to be facilitated by the right hippocampus. Assuming conserved lateralized function in rodents, it follows that both hippocampi are required for optimal performance of the water maze task (Fenton & Bures, 1993), but that their contributions to these tasks are different (Klur et al., 2009). Why then is the left hippocampus not needed for retrieval of a well-learned platform location as shown by Klur et al. (2009)? It may be that after extensive training to the point of asymptotic performance, memories acquired by the left hippocampus are consolidated into the cortex (Frankland & Bontempi, 2005), reducing or even abolishing the effects of left hippocampal inactivation. Additionally, why is the right hippocampus not needed for the Y-Maze spatial long-term memory task (Shipton et al., 2014)? In order to solve this task, mice were required to distinguish salient locations from each other, using distal spatial cues to identify which arm was rewarded. Once identified, their path to the goal location is restricted by the walls of the maze, and thus, does not require any route computation. Use of local cues, such as the maze walls, to guide navigation does not depend on the hippocampus (McDonald & White, 1994). Thus, as in humans, loss of left hippocampal function impairs spatial memory for salient places, such as a goal location. Loss of right hippocampal function impairs spatial navigation to well-learned places in rats (Klur et al., 2009) and to goal locations that the subject may even be constantly reminded of (Spiers et al., 2001).

### Short-term spatial memory in the left and right rodent hippocampus

Unilateral contributions to short-term spatial memory have also been tested in mice (Shipton et al., 2014). The left or right CA3 was inactivated during a spontaneous alternation task in the T-Maze, where mice will typically explore the arm they have not most recently visited if the inter-trial interval is brief. Mice started from the stem of the T-Maze and were allowed to choose one of the other arms to explore. Once they entered an arm, the entrance was closed, and they were allowed to explore only that arm for 30 seconds. After exploration, they were immediately placed back into the start arm and allowed to again choose an arm to explore. Under baseline conditions, mice will typically (∼80% of trials) choose to explore the arm other than the one they had most recently explored. Inactivation of either left or right CA3 impaired performance on this task, indicating a memory loss for the preceding exploration (Shipton et al., 2014). Interestingly, right inactivation led to even more impairment than left inactivation. In another test of short-term spatial memory, mice explored two arms of a three-arm Y-Maze during an encoding trial while a third arm was blocked off (Shipton et al., 2014). Mice were then returned to their home cage for one minute, unlike in the T-Maze task in which retrieval trials began immediately after the encoding trial ended. Mice were re-exposed to the apparatus and preference for the arm that was blocked off during encoding was measured. As in the T-Maze task, inactivation of either the left or right CA3 impaired preference for the novel arm in the Y maze. However, unlike in the T-Maze task, the impairments caused by left and right inactivation were equivalent. Although the authors did not remark upon this result in their conclusions, it is possible that the T-Maze alternation task was more impaired by right CA3 inactivation than by left CA3 inactivation because there was no delay between trials. Thus, it is possible that spatial working memory is right-dominant.

In summary, physiological and functional lateralization experiments suggest that left-lateralized synaptic plasticity enables the left CA3 to acquire and store spatial representations of particular locations and the events occurring at these locations as proposed by El-Gaby et al., (2015). Though not proposed in existing hypotheses of the bilateral hippocampus (discussed below), right-lateralized synaptic stability may enable spatial working memory and would always be required for spatial navigation.

## Previous Hypotheses of Hippocampal Lateralization and Spatial Memory

### Interhemispheric engram transfer

An early hypothesis of bilateral hippocampal function proposed that spatial memories are acquired by the left hippocampus and are then transferred to the right hippocampus for storage and retrieval (Klur et al., 2009). This is an elegant explanation for water maze impairments seen in rats following unilateral inactivation at different phases of acquisition and retrieval (Klur et al., 2009). Further, an experimental demonstration of memories translocating across hemispheres would profoundly change our understanding of how organisms acquire, store, and retrieve memories. However, this explanation does not appear to explain data from subsequent studies. First, split-brain mice without direct connections between the left and right hippocampi were capable of learning and retrieving a spatial Barnes Maze task (Shinohara et al., 2012). Second, long-term spatial memory on the Y-Maze task did not require the right hippocampus for storage or retrieval (Shipton et al., 2014; El-Gaby et al., 2016). Given these new data, it does not appear that hippocampal spatial memory engrams are transferred across hemispheres.

### The lateralized plasticity hypothesis

One promising hypothesis of bilateral hippocampal function suggests that left CA3 acquires and stores new memories via its capacity for synaptic plasticity and that right CA3 rapidly provides spatial representations of novel environments via stable neural networks that were preconfigured during development (El-Gaby et al., 2015). Prior to exploration of a novel environment, hippocampal ensembles have been shown to “preplay” compressed versions of the activity patterns that are later seen during spatial exploration (Dragoi & Tonegawa, 2011, 2014). It has been proposed that space can be represented rapidly by selection of established networks of hippocampal neurons (Dragoi & Tonegawa, 2014).

These networks may be a characteristic of right CA3 and would be preconfigured during development to represent a newly explored space rapidly upon initial exploration and these representations would be feedforward to bilateral CA1 (El-Gaby et al., 2015). Here, maps could be modified via spatial learning that engages left CA3, allowing for integration of spatial information acquired over many experiences to preconfigured spatial representations contributed by right CA3 (El-Gaby et al., 2015). Such modifications may establish new CA1 place cells that would integrate into networks with left CA3, reducing the contribution of the preconfigured right CA3 spatial representations to spatial memory over time (consistent with data reported by Shipton et al., 2014). The hypothesis proposed a right CA3 contribution to cognitive map formation. However, with spatial learning, right CA3 becomes dispensable. Thus, one limitation of the conceptualization by El-Gaby et al. (2015), is that it is not clear how it would account for the requirement of the right hippocampus when searching during the probe trial of a well-learned MWM (as seen in rats by Klur et al., 2009) or during spatial navigation (as seen in humans by Spiers et al., 2001).

## The DC Hypothesis: Discrete-Continuous Lateralized Spatial Representation

Here, I will propose a new hypothesis of bilateral hippocampal function that seeks to describe how the left and right hippocampi contribute to spatial memory and hippocampal function. Specifically, I concur with the previous proposal that left CA3 is specialized in associative memory acquisition and storage (El-Gaby et al., 2015), however, I add that right CA3 is *always needed* for route computation and navigation. This is in contrast to the proposal that right CA3 is less necessary for spatial memory with learning (El-Gaby et al., 2015). Thus, tasks that require retrieval of spatial memories will require left CA3 only if the stored memory has not been consolidated into the long-term neocortical store (e.g., “remote” memories), and will require right CA3 only if the task involves spatial navigation. Goal-oriented spatial navigation will engage both hemispheres (Maguire et al., 1998; Howard et al., 2014), although the processing in the left and right hemisphere during such a task will be distinct and complementary. Memories for particular locations (e.g., “start” and “goal” locations) are acquired, stored, and retrieved by left CA3 and routes between two such locations are computed by right CA3 (Figures 3 and 4). Thus, during goal-oriented navigation, recent place memories retrieved via left CA3 are held in a “goal working memory” network, while routes are computed and then updated in a right CA3 “spatial working memory” system. Further, I propose that these functions of left and right CA3 are accomplished by distinct ways in which they represent space: left CA3 represents space as discrete, salient locations, while right CA3 represents space continuously (Figure 3). These different types of spatial representation may facilitate the functions that have been reported to be lateralized in the hippocampus.

**Figure 3.**
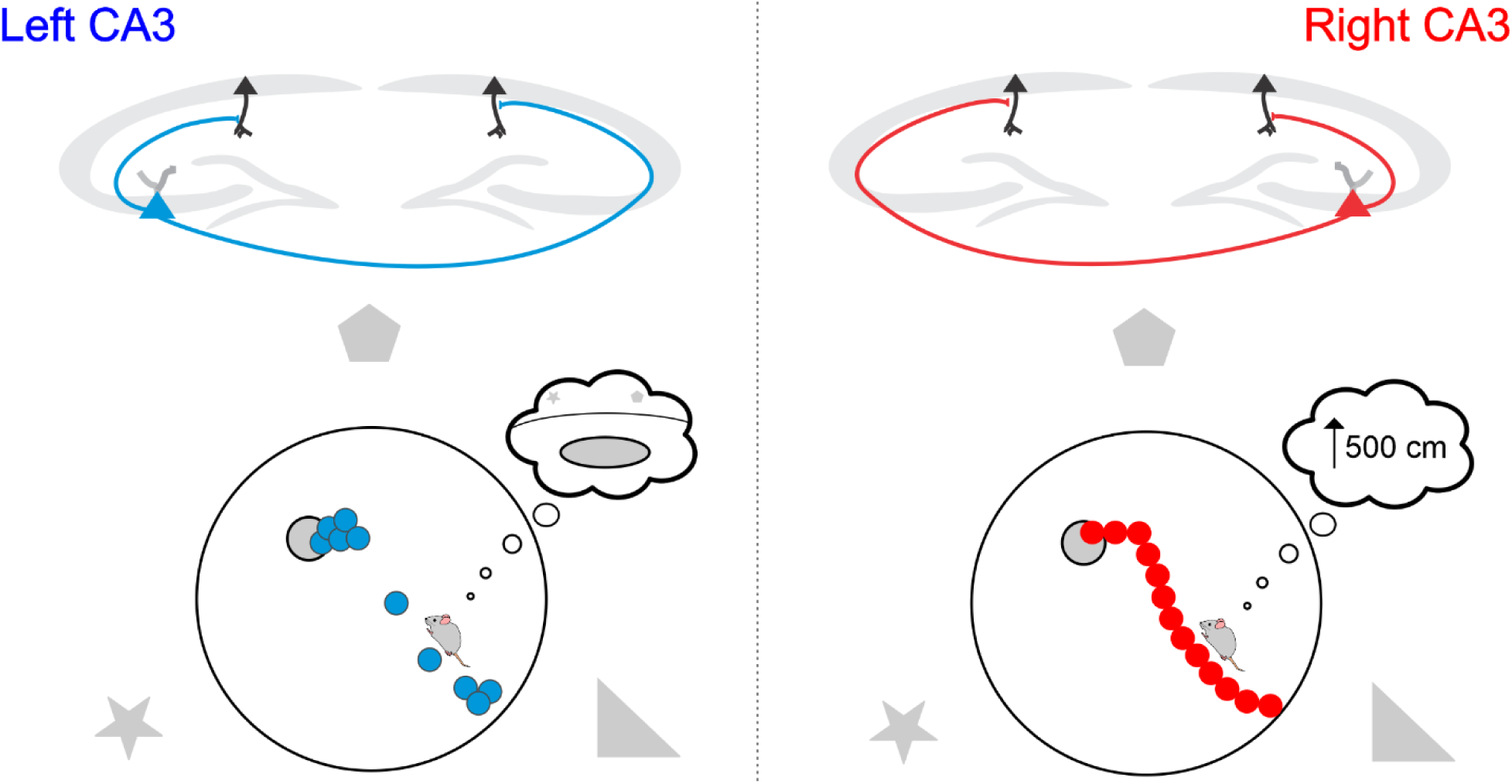
Lateralized discrete and continuous spatial representation in the hippocampus. Illustrated are the distributions of left (blue) and right (red) CA3 place fields of mouse during an MWM spatial navigation task according to the discrete-continuous hypothesis. In left CA3, spatial representation is non-uniform with greater weight allocated towards particularly salient locations on the map, such as an escape platform while non-salient locations along the traveled path are given less weight. Left-lateralized synaptic plasticity results in the storage and retrieval of salient locations as distinct places. In right CA3, spatial representation is uniform both at salient locations and at any given location along the traveled path. Right-lateralized synaptic stability allows for the computation of a navigational trajectory vector to the goal location. The illustration here is purely hypothetical and are not the depictions of any experimental data.

**Figure 4.**
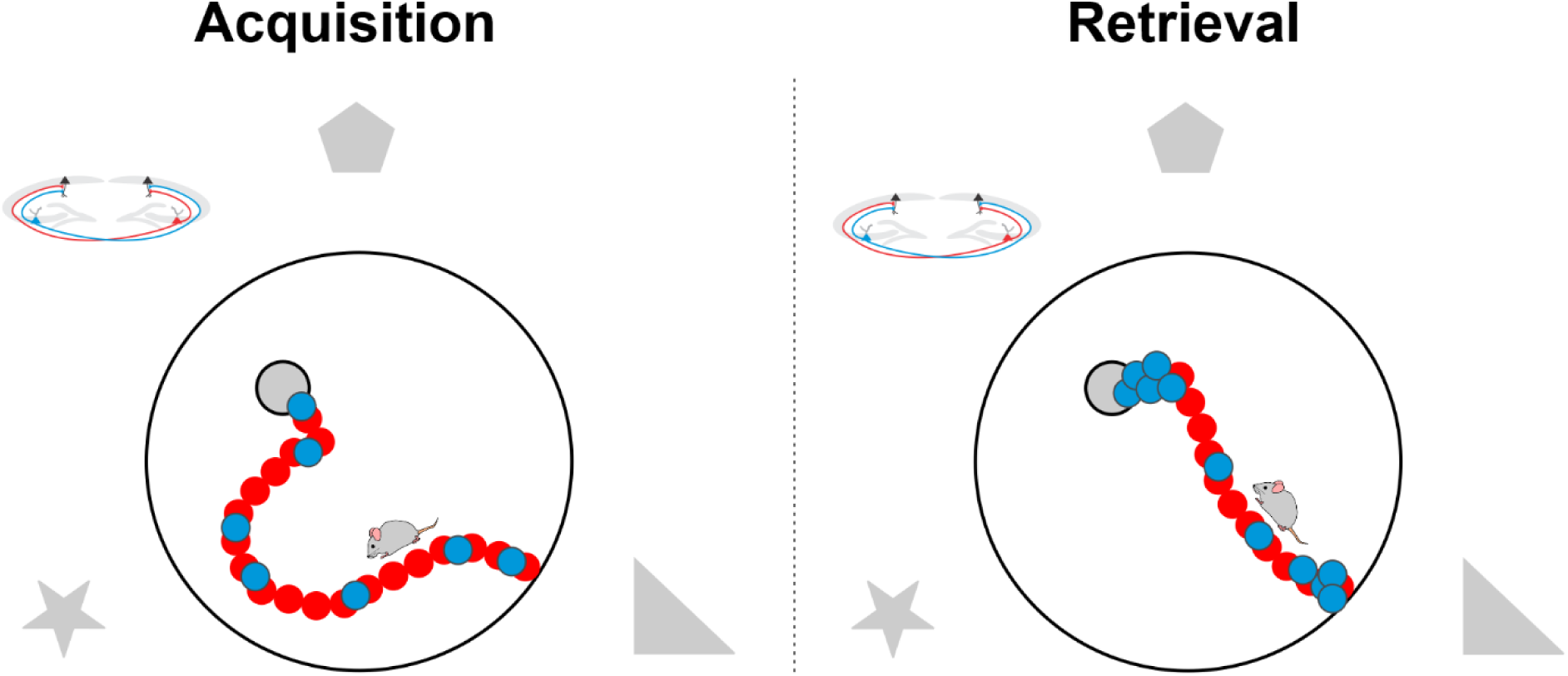
Lateralized spatial representations during acquisition and retrieval of a spatial memory in mice. Spatial representation of a water maze task in the hippocampus of mice. Upon first exposure to the water maze (Acquisition), left CA3 place fields (blue) are randomly and evenly distributed throughout the environment. However, learning-dependent synaptic plasticity leads to an accumulation of these place fields at discrete locations such as the start and goal locations, leading to non-uniform place field distribution in a well-learned maze (Retrieval). The formation and retention of these assemblies are required for later retrieval of the goal memory. During both acquisition and retrieval, right CA3 place fields (red) are distributed homogenously and are not changed by experience. These continuous representations allow for the computation of navigational distance and direction vectors. The illustration here is purely hypothetical and are not the depictions of any experimental data.

### Functional consequences of CA3 lateralization for spatial processing

Highly plastic networks in left CA3 may be specialized for acquisition and storage of new memories, and indeed a left hemisphere dominance of spatial memory acquisition has been demonstrated (Klur et al., 2009; Shipton et al., 2014; El-Gaby et al., 2016). Exactly what the function is of right-lateralized stability remains unknown. One hypothesis is that right CA3 was preconfigured during development and allows for rapid emergence of spatial maps when entering a new environment (El-Gaby et al., 2015). These maps can then be fed forward, establishing spatial maps in bilateral CA1 which are then modified over time by plastic projections from left CA3. My hypothesis extends this idea and proposes that stable networks in right CA3 are ideal for route computation, processing continuous spatial information (Figure 3) that is held in a right-lateralized spatial working memory system. This proposal is based on several findings of right hemisphere function in rodents an in humans. In particular, it is based on the finding that right CA3 inactivation produced a greater deficit than left inactivation in mice performing a short-term spatial T-Maze task in which retrieval trials immediately proceeded encoding trials (Shipton et al., 2014), a right-hemisphere dominance of route navigation in humans (Maguire et al., 1998; Spiers et al., 2001; Howard et al., 2014), and is consistent with rapid forgetting of spatial location after an interruption in humans following right temporal lobectomy (Smith & Milner, 1989). Thus, any task that would require retrieval of a memory stored in the hippocampus *in addition to* route-based spatial working memory would require both hippocampi.

Navigation to a specified goal location would depend on both memory of the goal location as well spatial working memory to guide navigation. Goal and other salient locations may be stored in left CA3 as associations between spatial cues. Further, these locations would be stored as discrete, individual places, preventing interference with other locations during retrieval (Figure 3). Thus, any task that requires the learning or retrieval of particular places within an environment or that requires discrimination of distinct places will be impaired by left CA3 inactivation. Indeed, optogenetic silencing of left CA3 impaired acquisition of the rewarded arm on a spatial long-term Y-Maze task in mice (Shipton et al., 2014; El-Gaby et al., 2016), pharmacological inactivation of the left hippocampus impaired acquisition of the spatial water maze task in rats (Klur et al., 2009), and left MTL damage impairs memory for specific places within a virtual environment in humans (Spiers et al., 2001). If salient locations are acquired, stored and retrieved by left CA3, why then does inactivation of left CA3 after acquisition not affect spatial memory in the MWM (Klur et al., 2009)? This may be because multiple training trials over six days to asymptotic performance resulted in the consolidation of the goal location engram into cortical memory networks (Frankland & Bontempi, 2005). Thus, only memories for individual locations that are not yet consolidated into the cortex would depend on the left hippocampus. This has not yet been explicitly tested. The short-term Y-Maze task is impaired by inactivation of left CA3 (Shipton et al., 2014), possibly because it requires encoding and retrieval of the three arms as discrete locations, allowing the mouse to determine the novel arm as a distinct location from the other two. Collectively, an assessment of left hemisphere loss-of-function studies supports the view that the left hippocampal system is specialized in memory for particular locations.

While space may be represented by left CA3 as discrete locations, it may be represented by right CA3 as continuous (Figure 3). Such continuous representations would be held in working memory, allowing for flexible navigation along routes. Any hippocampus-dependent task that would require spatial working memory, such as searching for a particular coordinate location within an environment, would be impaired by right CA3 inactivation. For instance, search for a well-learned location in the MWM is impaired by right hippocampal inactivation (Klur et al., 2009). Conversely, the *long-term* Y-Maze task, which does not require spatial working memory but only spatial memory for the reward location (discussed above), is not affected by right-inactivation (Shipton et al., 2014). Spatial working memory may be involved in the *short-term* Y-Maze task. As the animal explores the maze during the retrieval trial, a failure of spatial working memory may lead to less frequent re-entry to the novel arm, despite recognition of this arm as a distinct location from the two explored during the encoding trial. It is not surprising then that right CA3 in addition to left CA3 is needed to express novelty preference on the short-term Y-Maze task (Shipton et al., 2014). Humans with right MTL damage are impaired on navigation through a virtual environment, even when given a constant reminder of the goal location (Spiers et al., 2001). These data in humans and in rodents support the view that the right hippocampal system is specialized for route-based spatial navigation.

A curious case remains for the short-term T-Maze alternation task as this task is impaired by inactivation of either hemisphere but is more greatly impaired by right inactivation than by left (Shipton et al., 2014). Right CA3 inactivation led to near-chance levels of alternation on this task. As retrieval trials immediately followed encoding trials, and as I am proposing a right-hemisphere role in spatial working memory, this result is not surprising. However, left CA3 inactivation also impaired performance on this task, though not to the degree of right CA3 inactivation. Notably, when the encoding trial ended, the experimenter removed the mouse from its current arm and repositioned it in the start arm. Such a salient interruption may demarcate the encoding trial and retrieval trials as individual events, rather than maintaining them as a single, continuous event. Thus, just as I am proposing that the left CA3 may discretize space into salient locations, it may also discretize events into particular episodes, consistent with a left hippocampus dominance in episodic memory in humans (Spiers et al., 2001; Maguire & Frith, 2003). Thus, alternation on the T-Maze always requires spatial working memory (contributed by right CA3). Loss of left CA3 function impair may memory for the event that occurred during encoding, however, right CA3 may compensate for this loss somewhat, abating some of the deficit induced by the left CA3 inactivation.

### Spatial representation in left and right CA3

This framework of an interhemispheric difference in spatial representation as discrete (left) or continuous (right) parallels a framework proposed for explaining lateralization in the human visual system known as the categorical-coordinate hypothesis (Kosslyn, 1987 and 1994; for review, see Laeng et al., 2003). Behavioral, cognitive, and computational experiments have converged on the idea that in the human visual system, small, non-overlapping receptive fields in the left hemisphere discretize space and enable visual identification of spatial categories (e.g. on/off or above/below) while large, overlapping receptive fields in the right hemisphere represent space as a continuum and may be ideal for coordinate spatial representations, enabling one to determine how near or far two objects are from each other. Applied to the hippocampus, place cells in left CA3 may be small and non-overlapping, dividing space into discrete, salient locations where behaviorally relevant events are likely to occur. Place cells in right CA3 may be large and overlapping, providing continuity as one navigates through space. To date, there has not yet been evidence of hemispheric differences in hippocampal place fields. Many studies examine place fields unilaterally. Further, CA3 asymmetries may be lost in CA1 as this subfield receives input from both CA3. As place fields are most often recorded in CA1, a lack of CA1 place field lateralization may have discouraged investigation of place field lateralization in other areas. Chemogenetic inhibition of left or right CA3 in combination with calcium imaging of CA1 population activity will likely prove to be essential in determining left and right CA3 contributions to bilateral CA1 spatial mapping.

Although models of visuospatial lateralization have indicated how the spatial fields of individual cells can give rise to discrete and continuous representations, recent studies of spatial coding in the entorhinal-hippocampal system suggest that classic methods of excluding ‘non-spatial’ cells from analysis may be an insufficient approach to understanding spatial coding (Meshulam et al., 2017; Hardcastle et al., 2017). Instead, unbiased statistical approaches to cell classification has shown that both place and non-place cells contribute to place coding (Meshulam et al., 2017). It is possible that the lateralized coding motifs predicted above may be an antiquated view of hippocampal place coding. However, the underlying prediction that hippocampal activity patterns in the left hemisphere will code for particular locations while those in the right code for space continuously (Figure 3) remains unaffected. Goal locations (stored in left CA3) would be held in goal working memory during navigation and thus left CA3 neurons will often fire despite the organism’s location away from a discrete location stored in memory.

Therefore, using Bayesian decoders to understand population coding of the left and right CA3 will likely be critical to understanding their functions. During a spatial navigation task with a single goal, decoding of left CA3 activity would indicate a bias of representing the goal location, despite being far away from the goal at a given time (Figure 3). Tasks with multiple goals may cause left CA3 activity to cycle between the options until a decision is made, in which case activity would then be biased towards that goal location throughout navigation. Decoding right CA3 activity would likely accurately reflect the current position of animal and may reflect navigation choices to be made in the near future (Figure 3). Right CA3 position coding may be briefly modulated by left CA3 input if the animal has chosen a new goal location, initiating the computation of a new route.

Left and right CA3 representations would differ in how they are modified by experience. Discrete locations would be acquired by and stored in left CA3 via its capacity for synaptic plasticity. Thus, upon initial exposure to a novel environment, such as a water maze, left CA3 would show no bias in spatial representation (Figure 4). However, as learning proceeds, place fields would accumulate near salient locations, such as the escape platform. Although they are initially homogenous, left CA3 spatial representations will become skewed towards greater representation of discrete portions of space that the organism has learned are particularly important (Figure 4). Thus, left CA3 may be responsible for experience-dependent over-representation of goal locations reported in CA1 (Hollup et al., 2001; Gauthier & Tank, 2018) and inhibition of left CA3 during learning may prevent such changes to CA1 cognitive maps. Right CA3 will always homogenously represent space within an environment, regardless of how much learning about the environment has occurred (Figure 4). As these projections form anatomically mature and physiologically stable synapses with CA1 (Shinohara et al., 2008; Kohl et al., 2001; Shipton et al., 2014), inhibition of right CA3 is unlikely to affect learning-dependent changes in CA1. However, loss of continuous spatial information may impair CA1 coding of metric information required for navigation, such as direction and distance coding (Howard et al., 2014; Sarel et al., 2017). Modern *in vivo* 2-photon calcium imaging of neural activity allows for the recording of neural activity over large populations of hippocampal neurons in head-fixed awake behaving mice as they navigate through different virtual environments to find water (Dombeck et al., 2010). Spatial memory can be measured by a mouse’s lick rate near and away from water-rewarded locations (Lovett-Baron et al., 2014). These mouse virtual navigation tasks are reminiscent of virtual reality tasks used in humans that when combined with fMRI found hemispheric differences in hippocampal activity (Maguire et al., 1998) and can be used to test the predictions of the DC hypothesis in combination with invasive recording and inactivation methods. Recording calcium activity in CA3 during virtual navigation can determine if spatial representations in left CA3 are indeed weighted toward reward locations in an experience-dependent manner and whether right CA3 continuously represents space regardless of experience. The functional contributions of left and right CA3 to CA1 spatial coding and behavior can be tested using viral vectors to drive expression of inhibitory DREADDs receptors selectively in left or right CA3 (as done previously by Shipton et al., 2014 with inhibitory opsins) during 2-photon calcium imaging of CA1 neural activity to determine if left and right CA3 really do contribute to CA1 spatial navigation coding in distinct, but complementary manners. Inhibition of right CA3 during these tasks may impair accuracy of the search for the goal location, as evidenced by less licking at goal locations and more licking away from the goal. Inhibition of left CA3 may reduce licking specifically in the rewarded location, however, given the many trials required to train animals on such tasks, it is possible that goal location memory has been consolidated into the cortex and left CA3 inhibition may thus have no effect. Therefore, another behavioral strategy to determine whether left but not right CA3 is necessary for acquisition, storage and retrieval particular places is by testing left- or right-inhibited mice on a hippocampus-dependent contextual discrimination paradigm (Frankland et al., 1998). Left CA3 inhibition would lead to reduced discrimination between conditioned and alternative contexts. As mice are not required to navigate in such a task, inhibition of right CA3 would not be expected to have any effect. This paradigm can be easily adapted for use in other mammalian species such as a virtual context discrimination experiment in human subjects.

## Further Considerations

### Epigenetic lateralization

The left and right rodent hippocampi respond differently to experience. The effects of spatial learning on gene expression differ in left and right CA1 (Klur et al., 2009). After spatial training on the MWM, gene expression was heavily modulated in right CA1 (623 genes) but was modulated very little in left CA1 (74 genes), indicating epigenetic changes in response to spatial learning may be greater in the right hippocampus (Klur et al., 2009). Interestingly, spatial experience during early life was shown to produce epigenetic modifications in right, but not left CA1 (Tang et al., 2008; Shinohara et al., 2013). It will be important to identify the functions of genes that are upregulated or downregulated in one hemisphere and not the other following spatial learning. Doing so may lead to novel insights regarding the functions of the left and right hippocampi as well as to the broader question of how the hippocampus responds to experience.

### Is CA1 lateralized?

One major question remaining with respect to hippocampal lateralization is whether lateralization in function of CA3 extends to CA1? Though there are clear asymmetries at Schaffer synapses between CA3 axons and CA1 dendrites (Kawakami et al., 2003; Wu et al., 2005; Shinohara et al., 2008; Kohl et al., 2011; Shipton et al., 2014; Benito et al., 2016), both CA3 fields project bilaterally. Therefore, asymmetries in CA3 may not be relevant in CA1 as input from the left and right CA3 converge.

CA1 place cells are the most studied cell class of the hippocampus *in vivo*. To date, no form of lateralization has been reported with respect to place cell activity. However, circumstantial evidence indicates there may indeed be interhemispheric differences. One interesting debate in the hippocampal literature concerns how place cells represent an environment (O’Keefe, 2007). It is pointed out that many studies have reported that place fields are uniformly distributed across an environment (discussed in O’Keefe, 2007), while two studies have shown that they may actually cluster around salient cues or goal locations (Hetherington & Shapiro, 1997; Hollup et al., 2001). While some of the studies reporting uniform distribution do not report which hemisphere was recorded (O’Keefe, 1976; McNaughton et al., 1983; O’Keefe & Speakman, 1987), one study reported place fields in right CA1 to be distributed evenly across the recording to their chamber, and that the position or shape of place fields did not appear to be influenced by salient environmental cues (Muller et al., 1987). This was not the case in another study which found that place fields recorded in left CA1 appeared to cluster near salient environmental cues (Hetherington & Shapiro, 1997). Further, many place fields remapped following the removal of such cues. Similarly, place fields of left CA1 neurons have been shown to cluster around a goal location after learning, even when this goal was moved to a new location (Hollup et al., 2001).

Recently a new class of CA1 pyramidal neurons were described in the bat hippocampus, which were sensitive to either the distance or direction to a goal location (Sarel et al., 2017). Some of these goal-distance and goal-direction cells were shown to be distinct from place cells or head direction cells. Recordings were taken only from right CA1 (Sarel et al., 2017). Interestingly, goal-distance signals were originally reported in a human fMRI study where they occurred in the right, but not left hippocampus (Howard et al., 2014). Thus, if right-side specific, distance and direction cells may indicate lateralization in CA1.

### Does lateralization differ across sexes?

A majority of rodent hippocampal lateralization literature to date has only examined males. Sex hormones can have divergent organizational effects on spatial memory systems (Williams et al., 1990). It is not clear if there may be sex differences in the development of hemispheric differences in the hippocampus, although at this moment there is no data to indicate that this would be the case. In fact, if there is indeed a relationship between embryonic organization of organ laterality and hippocampal laterality (Kawakami et al., 2008; Goto et al., 2010), consistent organ laterality across sexes would indicate that consistent development of hippocampal laterality across sexes is likely. One study found no difference between adult males and female mice in left or right hippocampal c-Fos expression or in degree of lateralization (Jordan et al., 2019). While there is no data to suggest that hippocampal lateralization in males and females may be different, future studies will have to include females to determine whether or not an additional framework is needed (Shansky, 2019).

## Acknowledgments

I would like to thank Carolyn Pytte, Mohamady El-Gaby, Regis Shanley, Jeff Beeler, Dan McCloskey, Juan-Marcos Alarcón and André Fenton for useful discussions and comments on this manuscript. This work was funded by a Mina Rees Dissertation Fellowship.

